# Predictive Simulations of Gait with Exoskeletons that Alter Energetics

**DOI:** 10.1101/2021.08.31.458315

**Authors:** Anne D. Koelewijn, Jessica C. Selinger

**Affiliations:** Department Artificial Intelligence in Biomedical Engineering (AIBE), Faculty of Engineering, Friedrich-Alexander Universität Erlangen-Nürnberg, 91052 Erlangen, Germany; School of Kinesiology and Health Studies, Queen’s University, Kingston, ON K7L 3N6, Canada

**Keywords:** Biomechatronics, exoskeleton, gait simulation, metabolic rate, optimal control

## Abstract

Robotic exoskeletons, designed to augment human locomotion, have the potential to restore function in those with mobility impairments and enhance it in able-bodied individuals. However, optimally controlling these devices, to work in concert with complex and diverse human users, is a challenge. Accurate model simulations of the interaction between exoskeletons and walking humans may expedite the design process and improve control. Here, we use predictive gait simulations to investigate the effect of an exoskeleton that alters the energetic consequences of walking. To validate our approach, we re-created an past experimental paradigm where robotic exoskeletons were used to shift people’s energetically optimal step frequency to frequencies higher and lower than normally preferred. To match the experimental controller, we modelled a knee-worn exoskeleton that applied resistive torques that were either proportional or inversely proportional to step frequency—decreasing or increasing the energy optimal step frequency, respectively. We were able to replicate the experiment, finding higher and lower optimal step frequencies than in natural walking under each respective condition. Our simulated resistive torques and objective landscapes resembled the measured experimental resistive torque and energy landscapes. Individual muscle energetics revealed distinct coordination strategies consistent with each exoskeleton controller condition. Predicted step frequency and energetic outcomes were best achieved by increasing the number of virtual participants (varying whole-body anthropometrics), rather than number of muscle parameter sets (varying muscle anthropometrics). In future, our approach can be used to design controllers in advance of human testing, to help identify reasonable solution spaces or tailor design to individual users.

## 1 Introduction

This decade has seen rapid progress in the development of assistive devices designed to augment human locomotion. As early as the 1960s the first robotic exoskeleton prototypes, which are anthropometric in form and are designed to augment the user’s movements, were developed [1, 2]. However, early devices were often too heavy and cumbersome for practical use. Recent advances in both robotic hardware and software have pushed back technological boundaries, allowing for the development of more sophisticated, light weight, and intelligent designs [1–3]. Currently, a wide range of lower-exoskeletons are being tested in research labs around the world and even being sold commercially. Exoskeletons that augment the ankle, knee, hip, or some combination of joints exist. Some exoskeletons are active, or powered, using an actuator to apply torque to a joint, while others are passive, relying on springs or dampers to store and return energy during gait, without the need for an external power source. These devices can target walking and running gaits, as well as other activities of daily living. Potential applications of these evolving devices include: restoring bipedal locomotion for persons with paraplegia or persons post-stroke; enhancing the locomotor capabilities of military and emergency personal during load-carriage or complex terrain; reducing fatigue in unimpaired and older individuals; and providing robot-mediated physical therapy in a clinical setting [1–3].

Understanding how to optimally control exoskeletons during gait, to work in concert with a complex human user, is non-trivial. Given that humans tend to self-select gaits that minimize metabolic energy expenditure [4–11], a common high-level objective when designing and controlling exoskeletons is improved economy [3]. It may seem intuitive enough that an exoskeleton that mimics biological torques during gait should reduce the biological, or muscle generated, torque required at a given joint and in term whole-body energy expenditure. However, a number of complexities can disrupt this including muscles that cross multiple joints, elastic tissues that store and return energy in different phases of the gait cycle, complex interactions between the changing dynamics of the body and the adaptive strategies of the user, and individual user differences in both anatomy and neuromechanical control [12,13]. These factors perhaps explain why only in the last decade the ‘metabolic cost barrier’ was broken. That is, in 2013 Malcolm et al. used an exoskeleton, which applied assistive plantarflexion torques, to reduce the energy expenditure of able-bodied walkers below natural walking [14]. While many researchers had previously used heuristics or subjective feedback to select exoskeleton parameters, Malcom et al.’s approach included systematically testing a broad range of actuation profiles that varied actuation onset. They discovered that the optimal actuator onset timing, which reduced walking costs by 6% ± 2%, was in fact later than that predicted based on plantarflexor muscles at the ankle. While this brute parameter sweeping may be effective, its practically becomes limited with additional parameters and complexity of control, given the walking time and effort required to test each possible combination. Many powered exoskeletons have numerous tunable parameters at each joint, such as actuation timing, actuation magnitude, and stiffness, quickly resulting in a high dimensional optimization problem. To overcome this, ‘human-in-the-loop’ optimization methods have recently been developed, where a continuous measure of the user’s energy expenditure is fed into computer search algorithms that then iteratively adjust the device settings, in real-time, to minimize energy expenditure [15,16]. Zhang et al., using an evolutionary search algorithm that simultaneously optimized four exoskeleton parameters, demonstrated the largest energy savings to date—reducing the cost of walking by 24% ± 7% using a unilateral ankle exoskeleton [16]. However, like fine tuning based on subjective feedback or performance of full parameter sweeps, this method still relies on human testing, often necessitating hours of walking across multiple testing days.

Accurate and validated simulations of the interaction between exoskeleton devices and humans during gait have the potential to improve and expediate device design and control. Simulations can reduce the need for physical-prototyping of hardware, which can be costly and require substantial production time [17]. Beyond hardware, various control schemes and parameter settings for a particular device can be tested in simulation, reducing injury risk and time requirements for human experimental participants. Simulation even offers the ability to model individual differences and abilities, potentially helping to both extend findings and understand likely differences in clinical populations where the burden of experiential testing can be greatest [18]. Moreover, simulation can offer insight into measurements that can be difficult or impossible to capture during human testing, such as complex muscle-tendon interactions or individual muscle energy consumption. Of course, simulations are not without their own limitations that can hinder accurate predictions. While the anatomical detail of musculoskeletal models is ever improving, human anatomy is complex and cannot be fully characterized [19]. For example, modelling contact between the human foot and the ground is notoriously difficult and assumption laden [20], and muscle parameters are often based on limited in vitro experimental data [21]. We are also far from understanding the motor adaptation and learning processes that govern the human response to novel dynamics imposed by assistive devices. Despite these limits, we see an important role for simulation in the design and control of exoskeletons—one that is used in compliment with experimental testing to narrow the space of reasonable designs and control parameters.

Inverse dynamic simulations applied to musculoskeletal modelling has been used to investigate exoskeleton design and control. In this class of simulations, joint kinematics and total joint torques are often assumed to be unchanged from natural walking—in the simulation they are constrained to match experimentally measured data collected during walking without an exoskeleton [22–24]. Muscle generated joint torques can change as torques from the device are added, but total joint torques are fixed, as are joint angles. Solved outcome measures can include optimal device design or control parameters, the resulting individual muscle activations or energy expenditure, and the changes to muscle generated joint torques. For example, Van den Bogert (2003) used this approach to solve for the optimal geometric configuration of a lower-limb passive exotendon system that provided the greatest reduction in muscle generated joint torques [22]. However, an experimental study of this concept showed that while muscle generated joint torques decreased, joint angles changed and metabolic cost of walking was not reduced [25]. Another group has comprehensively investigated various actuated assistive device designs to reduce the metabolic cost of loaded walking and running [23, 24]. Their simulations demonstrated that frontal plane hip assistance may lead to the greatest improvements in economy and that ideal device torques can differ significantly from muscle generated torques in natural walking. However, while their simulations predicted large reductions in metabolic cost this was once again not reproduced in accompanying experiments [26]. The current discrepancies between inverse dynamic simulations and real-world human device testing may be because this approach constrains total joint torques and kinematics, failing to account for altered limb dynamics and the adaptive strategies of the user.

Forward dynamic simulations allow joint kinematics and muscle generated joint torques to change and may therefore overcome some of the limitations of inverse dynamic simulations when investigating exoskeletons. Furthermore, this approach allows for simulations of movements for which no experimental data is available, so-called predictive simulations. In this class of simulations, a high-level trajectory optimization problem is solved, where a periodic gait cycle is found that minimizes some objective function (or goal) [27–30]. The objective function often contains a number of weighted terms, such as maximizing smoothness, minimizing torques, and minimizing an energy term, be it muscular effort [27], metabolic energy [28] or both [29]. While these simulations can in principle be solved without requiring any experimental data, musculoskeletal model inaccuracies currently prevent many of these simulations from producing realistic gait without the use of reference data or expert input. Some researchers include a tracking term that weights kinematics and kinetics that are similar to references gait profiles [30], while others have expertly hand-tuned the objective function to produce a similar result [29]. Importantly though, these approaches do not fully constrain kinematics and kinetics, allowing adaptive strategies to emerge in response to the exoskeleton.

Methods to quickly and accurately solve predictive simulations of musculoskeletal models have only been developed in the last ten years [27], meaning their application to exoskeleton design and control has not been fully realized. Predictive simulations have been used to study other aspects of gait including, crouch gait in cerebral palsy [31], joint contact forces in running [32], shoe design on running efficiency [17], and loading asymmetry in prostheses [30]. A few groups have used predictive simulations investigate the optimal stiffness of passive ankle exoskeletons [18, 33]. However, these groups greatly simplified the musculoskeletal models, either limiting actuation to hip torques alone [33] or by not modeling tendon behavior [18]. These studies demonstrate that optimal gait kinematics and kinetics are likely to deviate from natural walking patterns in the presence of an assistive aid—both studies found this to be the case. However, accompanying experimental studies were not performed to validate these model predictions.

An existing limitation of predictive simulations is that they are often generated for a single set of musculoskeletal model parameters, limiting generalizability [18, 27–30, 33]. This is equivalent to generating predictions for a single participant, meaning conclusions may be dependent on the particular choice of anthropometric parameters. Recently, different approaches have been used to try to improve the generalizability of predictive simulations. Milller et al. (2013) produced simulations for four different participants, representing younger and older males and females [32]. Dorschky et al. (2019) pioneered the use of a set of ‘virtual participants’, where height and weight are randomly drawn from a reference set of participant data and, for each virtual participant, various sets of muscle parameters are randomly drawn from distributions that capture the variation expected in the population [17]. However, solving a large number of trajectory optimization problems in computationally costly and time consuming, and it is currently unclear how many virtual participants are required to make generalizable conclusions.

Our purpose in this study was to use predictive simulations of gait to investigate the effect of an exoskeleton that alters the energetic consequences of walking, using a virtual participant design. Specifically, we aimed to re-create a past experimental paradigm [4, 34], where robotic exoskeletons were used to shift people’s energetically optimal step frequency to frequencies higher and lower than normally preferred. In the human experiments, participants adapted their step frequency to converge on the new energetic optima within minutes and in response to relatively small savings in cost. To match the controller in this real-world experiment, we modeled a knee-worn exoskeleton that applied resistive torques that were either proportional or inversely proportional to step frequency—decreasing or increasing the energy optimal step frequency, respectively. We then performed predictive simulations of human walking with the device for a set of virtual participants, which we created to have similar anthropometric variables to our experimental participants. We compared our predictions to the experimental data to test if we could: i. replicate the exoskeleton torque applied at the knee joint during gait, ii. produce objective landscapes with optima at high and low step frequencies, and iii. solve for optimal gaits through adaptations in step frequency. We also investigated the sensitivity of our results to the sample size of virtual participants used and examined individual muscle energetics, offering insight into distinct coordination strategies adopted in response to the exoskeleton.

## 2 Methods

### 2.1 Musculoskeletal Model

We used a sagittal plane nine-degree of freedom musculoskeletal model of the lower limb (Fig. 1). The model consists of seven segments (a trunk, two upper legs, two lower legs, and two feet) connected via revolute joints. The degrees of freedom are the position and orientation of the trunk, flexion-extension of each hip, flexion-extension of each knee, and dorsiflexion-plantarflexion of each ankle. The model is actuated by eight muscles in each leg, including: iliopsoas, gluteals, quadriceps, hamstrings, vastus, gastrocnemius, soleus, and tibialis anterior. The muscles are modeled as three element Hill-type muscles with a series elastic element, parallel elastic element, and a contractile element with activation dynamics, force-length, and force-velocity properties. We calculated individual muscle metabolic rate using the model by Margaria [35]. Ground contact was modelled with a penetration-based model [30]. The multibody dynamics and ground contact model were derived using Autolev (OnLine Dynamics, Inc., Sunnyvale, CA, USA), combined with muscle dynamics coded in C, and compiled as a MEX-function [27, 30].

**Figure 1:**
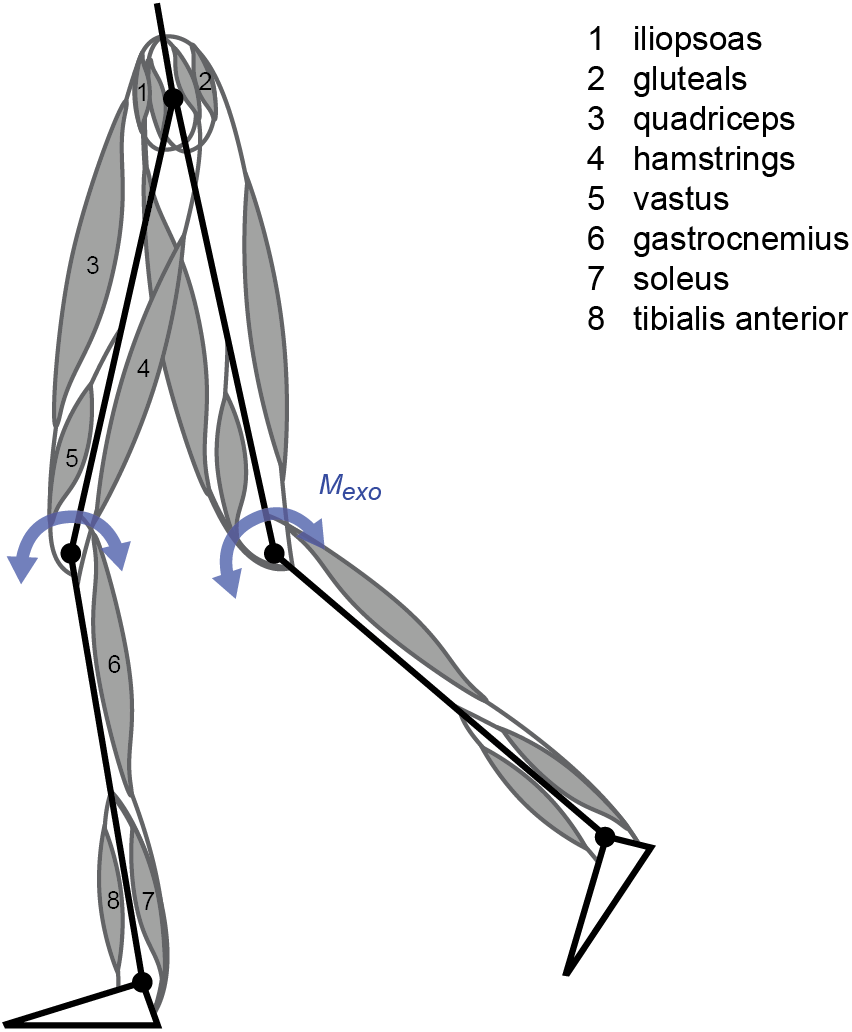
Musculoskeletal model and exoskeleton torque. Our nine degree of freedom model includes eight muscles at each leg (1-8) and an exoskeleton flexion-extension torque added to each knee (curved arrows).

### 2.2 Exoskeleton Model

We then added an ideal, massless, exoskeleton to our simulations by applying torques at the knee joints that resisted both knee flexion and extension, and thereby added an energetic penalty. We applied two different exoskeleton controller conditions: *penalize-high*, where higher step frequencies were penalized such that the optimal step frequency was lower than natural, and *penalize-low*, where lower step frequencies were penalized such that the optimal step frequency was higher than natural. We designed these controllers to replicate our real-world exoskeleton controllers [4].

To accomplish this, we made the *applied torque* (*M*_*exo*_) proportional to *step frequency* (*s*) throughout the stride and proportional to knee *angular velocity* 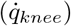 within the stride. To ensure that the exoskeleton torque was penalizing, we determined its absolute value and multiplied this by the opposite sign of the knee *angular acceleration* 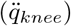. The absolute peak torque was limited to 12 Nm. This yields the following equation:

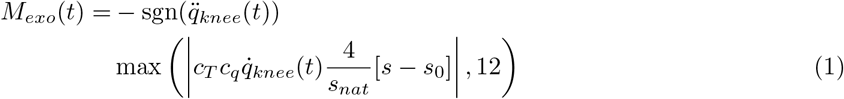

where *t* is time, *c*_*T*_ = 3.36×10^*−*2^ Nm/A is the motor’s torque constant and *c*_*q*_ = 60 C/rad is a constant that represents the relationship between motor current (A) and knee angular velocity (rad/s). The *zero-torque step frequency (s*_0_) is set to −15% of the *natural step frequency* (*s*_*nat*_) in the penalize-high condition and +15% of the natural step frequency in the penalize-low condition.

To allow the optimization to be solved with a gradient-based algorithm, we converted Equation 1 to a twice-differentiable function. This involves three steps.

First, the absolute value of the applied torque (*M*_*exo,abs*_) was determined using the following soft absolute function:

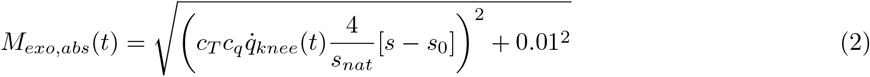

Second, the applied torque was limited to an absolute peak of 12 Nm (*M*_*exo,lim*_) using an exponential soft-max function:

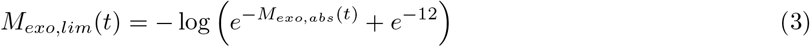

Third, the sign function was approximated using the inverse tangent:

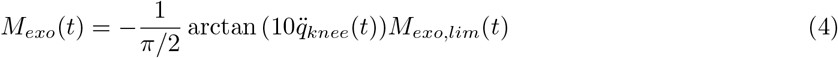

where we used the factor 10 to increase the nonlinearity and similarity to the sign function.

### 2.3 Trajectory Optimization Problems

We generated muscle-driven simulations of walking at 1.3 m/s by solving trajectory optimization problems. The objective was to minimize: i) a weighted sum of muscular effort, which was calculated as the cubed muscle stimulation; ii) a tracking error, which was calculated as the difference between predicted and literature joint angles and ground reaction forces during normal walking [36]; and iii) a regularization term [28, 37]. The weighting ratio between effort and tracking error was 1000:1, a factor one hundred times higher than previous work with the same model [30]. We chose this high ratio to allow simulations to deviate from tracked data as much as possible, and therefore adapt step frequency, while still maintaining realistic gaits. We included a small regularization term to minimize the derivatives of the states and controls, which enhances convergence of the optimization without affecting simulation accuracy [28, 37]. We assumed symmetry and simulated only half a gait cycle. This yields the following optimization problem:

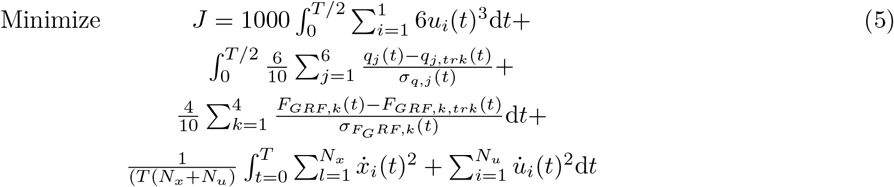

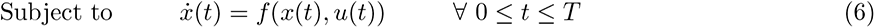

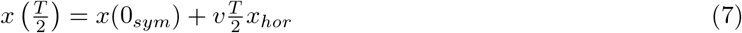

where *T* is the duration of the gait cycle, *u*_*i*_ is the stimulation of muscle *i, q*_*j*_ is the angle for joint *j, q*_*j,trk*_ is the tracked angle for joint *j* [36], *F*_*GRF,k*_ is the ground reaction force signal *k* for the left and right legs in vertical and horizontal direction, *F*_*GRF,k,trk*_ is the tracked ground reaction force for signal *k, σ* is the standard deviation of the tracked variables, *v* is the walking speed, *N*_*x*_ is the number of states, and *N*_*u*_ is the number of muscles.

Equation 6 describes the dynamics, which are dependent on the state 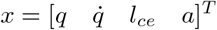, with contractile element length *l*_*ce*_ and muscle activation *a*, and the input (muscle stimulation, *u*). Equation 7 denotes the periodicity constraint, the subscript *sym* denotes the mirror of the state, meaning that the joint angles and angular velocities were switched between the left and right leg, and the subscript *hor* indicates the states which should translate horizontally.

We created walking simulations for three conditions: *natural*, where no added knee torques were applied by the exoskeleton, as well as *penalize-high* and *penalize-low*, where each respective exoskeleton controller applied torques to the knees. We added the applied exoskeleton torques to the torques generated by the muscle forces. For each condition, we first optimized step frequency to investigate if we could predict the energy optimal gait adaptation. This optimal step frequency is the step frequency where the full objective (Equation 5) is minimized. Next, we created simulations at a range of fixed step frequencies (−15% to +15% of the natural optimal step frequency, at increments of 1%) to generate the *landscapes* for all three conditions (natural, penalize-high and penalize-low). We created a *full objective landscape*, which is the full objective function evaluated for the range of fixed step frequencies, an *effort term landscape*, which is the effort term of the objective function alone evaluated for the range of fixed step frequencies, and the *metabolic rate landscape*, which is a sum of the individual muscle metabolic rates (to estimate whole-body metabolic rate) for the range of fixed step frequencies. The step frequency was fixed by constraining the duration of the gait cycle (*T* = *T*_*fixed*_).

To create all simulations, we solved trajectory optimization problems with direct collocation and a backward Euler discretization. We used 30 nodes per half gait cycle. We coded the problems in MATLAB R2018a (Mathworks, NA, USA) and used IPOPT [38] to solve the resulting large-scale constrained optimization problem. We first created a simulation of standing by constraining the derivatives of the degrees of freedom to be equal to zero: 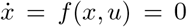 while minimizing muscular effort, as described in [37]. We repeated this problem with 50 different random initial guesses and used the solution with the lowest objective. Then, we used standing as an initial guess to find walking simulations with free, or optimal, step frequencies in the natural, penalize-high, and penalize-low conditions. Next, to create the landscapes, for each condition we first solve for a simulation where the step frequency is equal to the optimal step frequency in the natural condition. To find these simulations for each condition, we use the previously solved walking simulations with free, or optimal, step frequencies for the same condition as an initial guess. Next, for each condition, we used the simulation with step frequency fixed to the natural optimum as an initial guess for simulations fixed at +1% and −1% of the natural optimum. For each condition, we then used these solved simulations as initial guesses to find the simulations fixed at +2% and −2% of the natural optimum, and we repeated this process until we reached +15% and −15% of the natural optimum, the full range of step frequencies for the landscapes [4].

### 2.4 Virtual Participant Study

To simulate natural variation between individuals, we created a set of 50 virtual participants. We generated *anthropometric parameters* for each virtual participant by drawing from distributions of mass and body-mass index (BMI). We set these distributions to be the same as that measured for our experimental participants: 65.3 ± 9.8 kg (mean ± SD) for mass and 23.3 ± 2.3 kg/m^2^ (mean ± SD) for BMI [4]. We used mass and BMI, instead of mass and height, since mass and height are dependent variables. Within each virtual participant we also produce variation in *muscle parameters* to account for uncertainty by generating 10 different sets of muscle parameters. To do so, we first scaled the maximum isometric force according to the body mass. Then, we drew the maximum isometric force, optimal fiber length, muscle-tendon length, width of the force-length curve, maximum shortening velocity, maximum force during lengthening, Hill parameter, muscle moment arms, stiffness of the parallel elastic element, strain in the series elastic element, and the activation and deactivation time constant from normal distributions with the means equal to commonly used nominal values and the standard deviations equal to 10% of nominal [17, 21, 39]. Finally, we set the series elastic element slack length such that the muscle tendon length in neutral skeleton pose (where all joint angles are zero) was equal to the nominal value [17]. In total, this yields 48,000 simulations (1 simulation at a free, or optimal, step frequency + 31 simulations at various fixed step frequencies for the energy landscape × 3 conditions = 96 simulations; 96 simulations ×10 muscle parameter variations × 50 virtual participants = 48,000).

### 2.5 Analysis

We first identified and removed all simulations of virtual participant and muscle parameter set combinations that produced at least one unrealistic simulation. We identified simulations as unrealistic when the difference in metabolic rate between at least one simulation with fixed step frequency and free step frequency in the same condition was larger than 2 W/kg. We further verified these outliers by visual inspection of the gait cycle, to confirm that the gait was unrealistic and likely the result of a local minimum. For all simulated outcome measures, we then averaged across the remaining muscle parameter variations within each virtual participant. We then calculated the mean and standard deviation of the outcome measures across all 50 virtual participants. We compared the average height and body mass of our virtual and experimental participants to ensure they were similar.

To investigate the quality of predictions from our simulations, we compared the simulation results to the experimental results of Selinger et al. (2015) [4]. Simon Fraser University’s Office of Research Ethics approved the protocol, and participants gave their written, informed consent before experimentation. We determined the coefficient of determination (R^2^), between simulation predictions and experimental data, for the across-stride average and within-stride torque profiles in the penalize-high and penalize-low conditions, as well as for the objective, effort, and energy landscapes in all three conditions. To compare the optimal, or freely chosen, step frequencies and the accompanying full objective, effort term, or metabolic rate values between the experimental and simulated data, we calculated the percent changes from the natural condition to the penalize-high and penalize-low conditions. We then compared the means and standard deviations of theses percent changes between the experimental and simulated data.

We also investigated the changes in simulated metabolic rate for individual muscles across the three conditions. For all three conditions, we generated and compared *individual muscle metabolic rate landscapes*, which were the individual muscle metabolic rates computed for the range of fixed step frequencies. For simulations with an optimal, or freely chosen step frequency, we also determined the percent change in individual muscle metabolic rate between the natural and the penalize-high and the penalize-low conditions, during both stance and swing phase. To split individual muscle metabolic rates throughout the stride into stance and swing, we used the vertical ground reaction force to determine when the foot was in contact with the ground. To calculate the relative contribution of stance and swing to metabolic rate, we divided by the full duration of the gait cycle, such that the sum of both was equal to the total metabolic rate.

Finally, we investigated how different virtual participant and muscle parameter set sample sizes affected our simulation outcomes. We repeated our previously described analysis to determine the percent change in optimal, or freely chosen, step frequencies between the natural condition and the penalize-high and penalize-low condition for virtual participant sample sizes between 1 and 50 and muscle parameter sample sizes between 1 and 10. We also calculated the accompanying percent change in whole-body metabolic rate. To accomplish this, we start with the full set of 50 virtual participants by 10 muscle parameters sets. For each of the desired number of virtual participants and muscle parameter sets, we randomly select samples from the full set. We then repeat this sampling 10 times, to account for randomness in the data, and then calculate the percent difference between the outcome measures of the sample and the full dataset. For each outcome measure, we then calculate the root mean square (RMS) error across these 10 repeats for each combination of sample sizes.

## 3 Results

Of the 500 combinations of virtual participants and muscle parameter sets (50 × 10), we removed 6 that produced unrealistic simulations. Our remaining virtual participants had anthropometrics similar to the experimental participants. Virtual and experimental participants had an average height of 1.64 ± 0.16 m (mean ± SD) and 1.67 ± 0.10 m, respectively, while average body masses were of 63.9 ± 10 kg and 65.3 ± 9.8 kg, respectively. The mean time required to solve all 96 gait simulations for one virtual participant with one muscle parameter set was 57 ± 30 minutes, ranging between 35 and 705 minutes. The simulations were solved on a Fujitsu Celsius M740 workstation with a Xeon E5-16xx processor with an Ubuntu 18.04.5 LTS operating system.

### 3.1 Comparison of experimental and simulated outcome measures

Our simulated exoskeleton torques within the stride and averaged across the stride, resembled that from human experiments (Fig. 2). The coefficients of determination between the simulated and experimental within-stride torques ranged between 0.33 and 0.49. For both the penalize-high and penalize-low controllers, our simulated within-stride torques better matched experimental when angular velocities were greater, during late stance and swing (*>*50% of gait cycle, Fig. 2AB). During early stance within-stride estimates were poorer, though still exhibited two moderate peaks. Although within-stride differences existed in the torque profiles, the average simulated torques across strides were very consistent with that applied in human experiments (Fig. 2C). The coefficients of determination between the simulated and experimental across-stride torques were 0.97 and 0.98 for the penalize-high and penalize-low conditions, respectively. The penalize-low and penalize-high conditions exhibited the desired relationships with step frequency: being negatively and positively sloped, respectfully, and delivering the same magnitude torque at the 0% step frequency.

**Figure 2:**
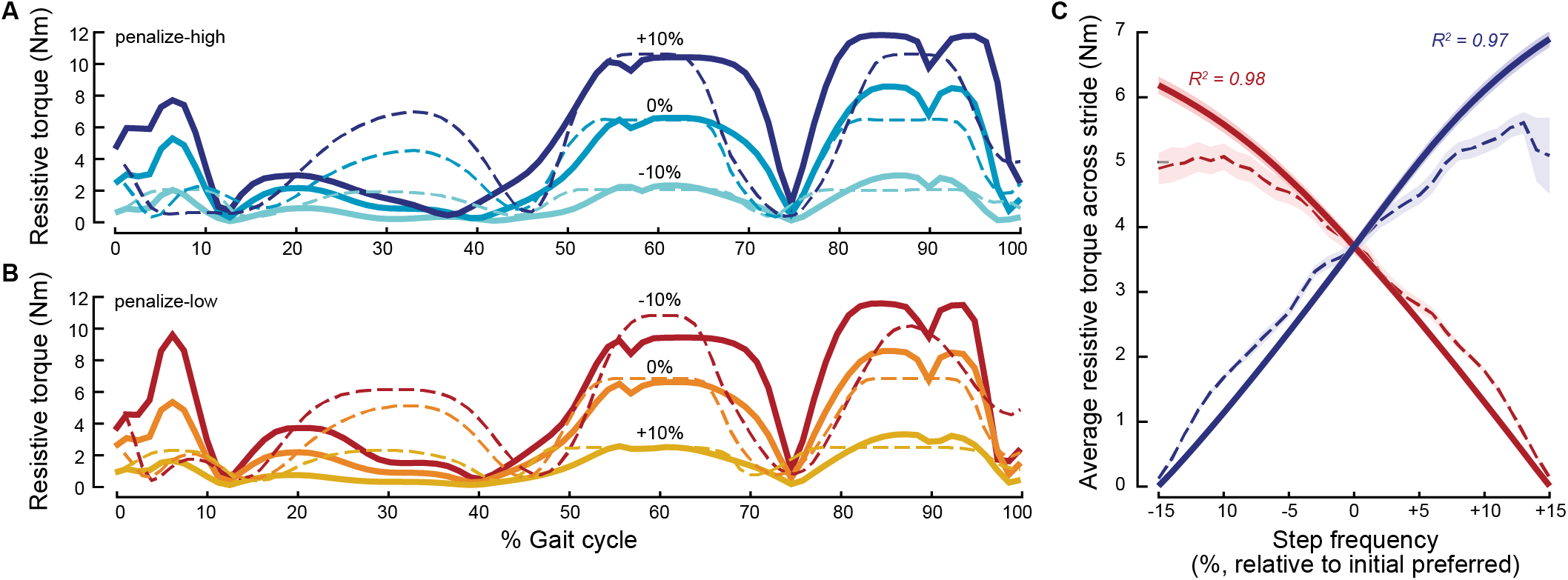
Comparison of simulated and experimental within-stride and across-stride torques. Average torques throughout the gait cycle for the penalize-high condition (A) and penalize-low condition (B) at −10%, 0% and +10% of natural step frequency. C. Average torques across the stride for the penalize-high (blue) and penalize-low (red) conditions.

The torques applied by our simulated exoskeleton controllers produced full objective, effort term, and metabolic rate landscapes similar in shape to human experimental energy landscapes (Fig. 3), though differences existed in the non-normalized natural optimal step frequencies and metabolic rates. In all landscapes, the simulated penalize-low controller produced a negatively sloped gradient about the initial preferred step frequency (0%) and shifted the optimal gaits to higher step frequencies, as was the case in the experimental energy landscapes. The simulated penalize-high controller produced a positively sloped gradient about the initial preferred step frequency (0%) and shifted the simulated energy optimal gaits to lower step frequencies, once again in a manner consistent with the experimental landscapes. However, in the full objective landscape the optimal step frequency occurred at 2.24 Hz, which is higher than the step frequency of the tracked data and the experimental optima, which were both 1.8 Hz [4,36]. The effort term and metabolic rate landscapes in the natural condition had optimal step frequencies that were both lower than that for the full objective landscape (2.14 Hz and 2.16 Hz, respectively), but were still high. Moreover, the simulated energy landscapes predicted metabolic rates that were notably higher than the experimental metabolic rates (+0.5 to 1.5 W/kg), as well as smaller relative changes in metabolic rate under the penalize-high and penalize-low conditions. Overall, the coefficients of determination, between the simulated and experimental landscapes, ranged between 0.62 and 0.98 and coefficients tended to be higher for the penalize-high and penalize-low conditions than the natural condition.

**Figure 3:**
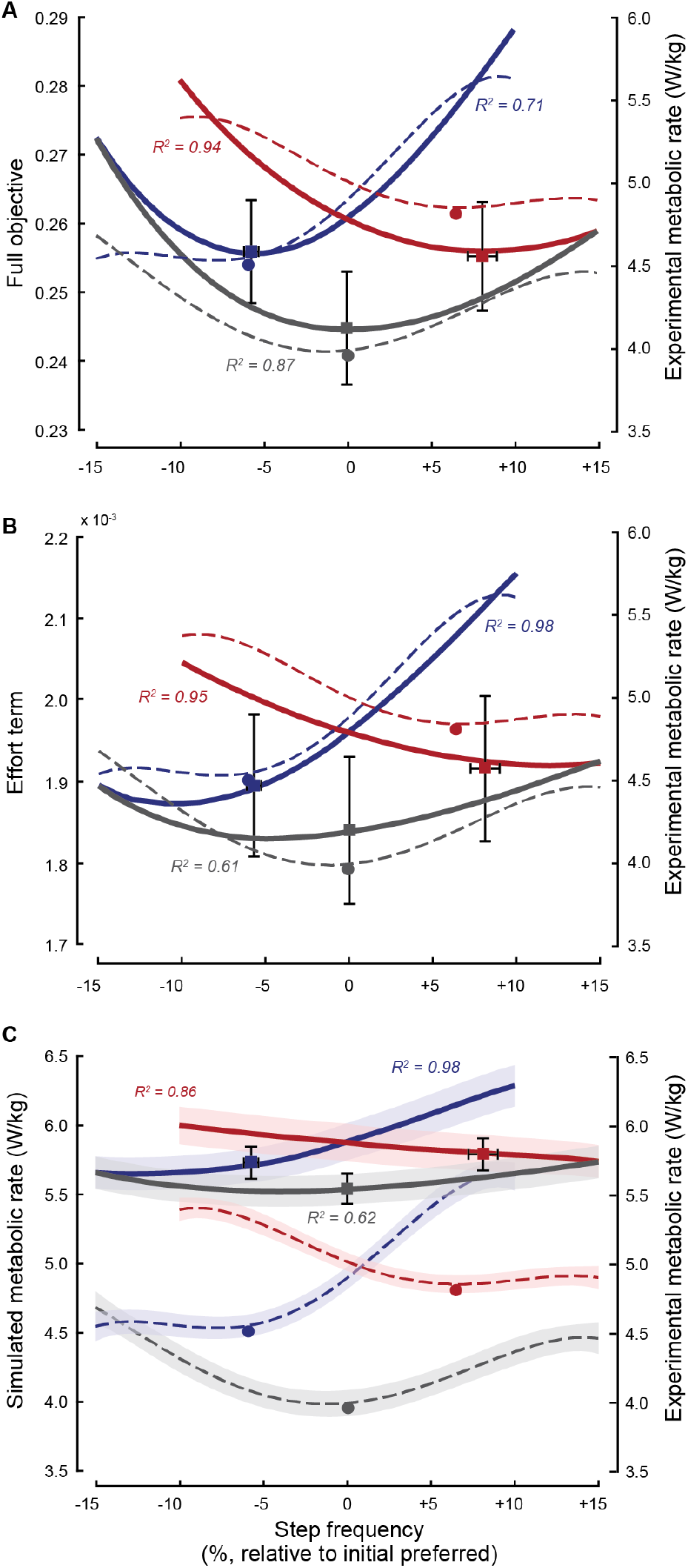
Comparison of simulated full objective, effort term, and metabolic rate landscapes with experimental metabolic rate landscapes. Experimental metabolic rate landscapes as well as simulated full objective (A), effort term (B), and metabolic rate (C) landscapes for the natural (grey), penalize-high (blue), and penalize-low (red) conditions. In all plots, solid lines are simulated data, while dashed lines are experimental data [4]. Squares indicate optimal gaits, that minimize the full objective, for each simulated condition, with error bars representing the standard error, while the circles indicate experimental optimal gaits for each condition. In C, shading represents standard error.

Like our human participants, our virtual participants responded to the new landscapes, adapting toward the energy optimal step frequencies (Fig. 3). Under the penalize-high and penalize-low conditions virtual participants adapted their preferred step frequency by −5.7% ± 1.5% and 8.1% ± 3.1%, respectively (these step frequencies are those that minimize the full objective). These adaptation magnitudes were both indistinguishable from those displayed by experimental participants in each respective condition (−5.7% ± 3.9%, p =0.79; −6.9% ± 4.3%, p =0.16). In the penalize-high condition, virtual participants’ full objective magnitude at the new preferred step frequency was 1.9% ± 1.7% lower than the full objective magnitude at the initial preferred step frequency under the penalize-high control function (Fig. 3A). For the effort term the equivalent reduction was 3.6% ± 2.4% (Fig. 3B), while for the simulated metabolic rate the reduction was 2.5% ± 0.91% (Fig. 3C). These reductions are smaller than the 8.1% ± 7.0% reductions in metabolic rate seen in the experiment under the penalize-high condition (Fig. 3C, [4]). In the penalize-low condition, virtual participants’ full objective magnitude at the new preferred step frequency was 2.2% ± 2.1% lower than the full objective magnitude at the initial preferred step frequency under the penalize-low control function (Fig. 3A). For the effort term the equivalent reduction was 2.5% ± 2.5% (Fig. 3B), while for the simulated metabolic rate the reduction was 1.4% ± 1.0% (Fig. 3C). These reductions are again smaller than the 4.0% ± 3.8% reductions in metabolic rate seen in the experiment under the penalize-low condition (Fig. 3C, [4]). Similar to the experiment, the simulation predicts that the changes in effort and metabolic rate in the penalize-low condition are roughly double those changes in the penalize-high condition, but this was not observed for the full objective.

### 3.2 Effects on Individual Muscle Metabolic Rate

Individual muscle changes in metabolic rate, across the energy landscapes, offer insight how the exoskeleton controllers produce changes in whole-body metabolic rate. Across the full landscape (−15% to +15% change in step frequency) the range of metabolic rates (difference between the minimum and maximum rate across penalize-high, penalize-low, and natural) are similar, about 0.3 W/kg, for all muscles except for the vastus, where the range is 0.6 W/kg (Fig. 4). This indicates that the vastus has the largest influence on the wholebody energy landscape, while all other muscles have a similar, but lower, influence. We also found that individual muscle changes in metabolic rate across the energy landscape differed between muscles that cross the knee and those that do not. Under natural conditions, muscles that cross that knee (hamstrings, rectus femoris, vastus, and gastrocnemius) tend to display an optimum (minimum metabolic rate) within the range of step frequencies in the landscape, while those muscles that do not cross the knee (iliopsoas, gluteals, soleus, and tibialis anterior) tend to have a consistently increasing or decreasing slope across the landscape. Although the steepness of these slopes is altered by the exoskeleton controller conditions, the direction remains unchanged (i.e iliopsoas has a positively sloped gradient for natural, penalize-high and penalize-low conditions). Conversely, for muscles that cross the knee, the two exoskeleton controllers create slopes of opposite direction—creating a positively sloped gradient under the penalize-high condition (decreasing the optimal step frequency) and a negatively sloped gradient under the penalize-low condition (increasing the optimal step frequency). For these muscles, the penalize-high condition tends to create steeper slopes than the penalize-low condition, the effect of which is evident in the whole-body metabolic rate landscapes.

**Figure 4:**
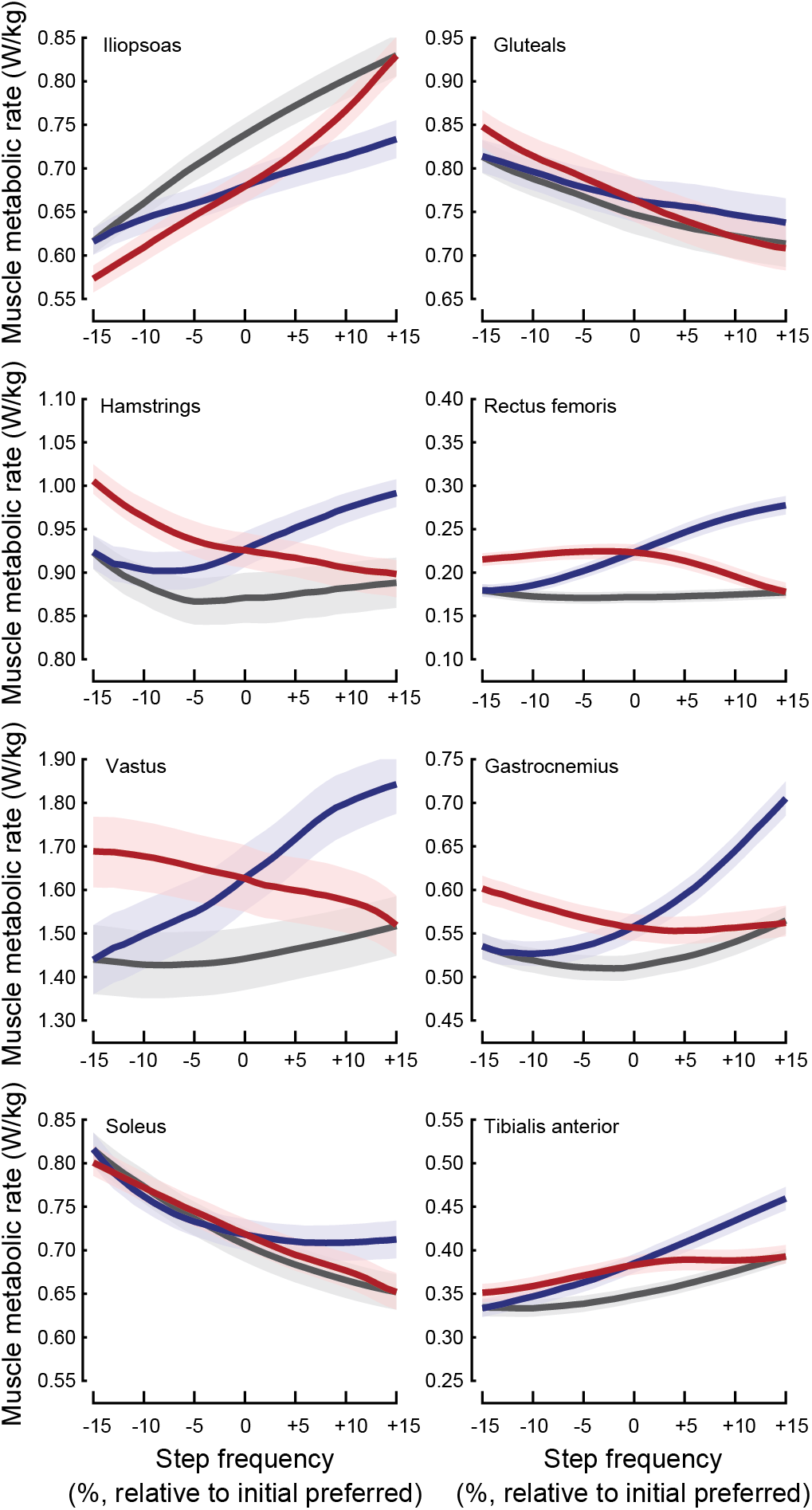
Simulated individual muscle contributions to energy landscapes. Simulated muscle metabolic rate across step frequency for the penalize-high (blue), penalize-low (red) and natural (black) conditions. In all plots, solid lines show the average metabolic rate, across 50 virtual participants, while the faded fills show the standard error. All muscle y-axes span the same magnitude range (0.03 W/kg), accept the vastus, which spans 0.6 W/kg.

An examination of simulated metabolic rate at the individual muscle level during the stance and swing phase reveals distinct coordination strategies consistent with each exoskeleton controller condition (Fig. 5). Our simulated optimal gaits show changes in muscle metabolic rate not only for muscles spanning the knee, but also muscles crossing the hip or ankle. In the penalize-high condition, muscle metabolic rate decreased across more swing muscles, but showed large increases for muscles crossing the ankle during stance (i.e. iliopsoas, swing: −9%; tibialis anterior, stance: +15%). This is consistent with an adaptation toward lower step frequencies, requiring more work during push off at the ankle to lengthen steps and relatively less work to swing the limb. In the penalize-low condition, the opposite was the case; muscle metabolic rate increased across nearly all muscles during swing, but showed decreases for many muscles during stance, particularly the tibialis anterior that crosses the ankle (i.e. gluteals, swing: +35%; tibialis anterior, stance: −21%). This is consistent with an adaptation toward higher step frequencies, requiring more work at the hip during swing and relatively less work during a shorter stance phase.

**Figure 5:**
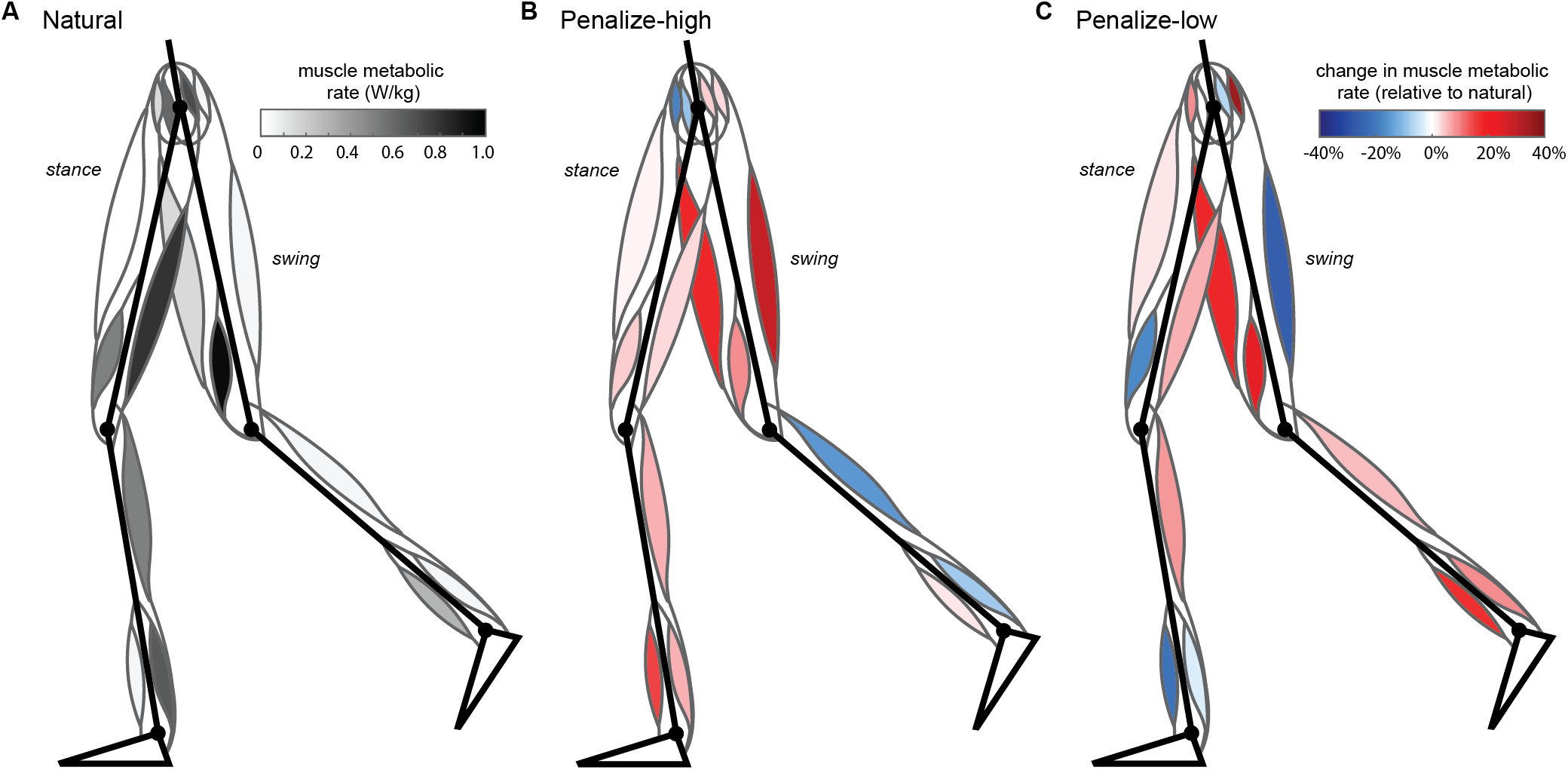
Effects of exoskeleton on simulated individual muscle metabolic rate. Average muscle metabolic rate, during stance (left, standing leg) and swing phase (right, swinging leg), in W/kg for the natural condition (A), and relative to natural for the penalize-high (B) and penalize-low (C) conditions.

### 3.3 Virtual Participant Study

We found that both simulated step frequency adaptations and metabolic rate were primarily affected by the number of virtual participants, although increasing the number of muscle parameters sets had an effect at larger virtual participant numbers (Fig. 6). The RMS difference, between outcomes with a limited dataset and that with the full dataset (50 virtual participants and 10 muscle parameter sets), clearly increases when the number of virtual participants is decreased, while there is a smaller effect when the number of muscle parameter sets is decreased, for both step frequency (Fig. 6A) and metabolic rate (Fig. 6B). For example, when using only one virtual participant and one set of muscle parameters, the RMS difference in step frequency is 1.9% in the penalize-high condition and 3.8% in the penalize-low condition. These differences are high, being roughly half the predicted step frequency adaptation magnitudes under the full dataset. However, when using all 50 virtual participants and one set of muscle parameters, the RMS difference decreases to less than 0.2% in the penalize-high condition and less than 0.4% in the penalize-low condition (a ten-fold reduction). Conversely, if only a single virtual participant is used and all 10 sets of muscle parameters are included, the RMS difference remains high, at 1.5% in the penalize-high condition and 3.0% in the penalize-low condition. Similar results were observed for the metabolic rate. However, it does become more advantageous to increase the muscle parameter set size, instead of the number of virtual participants, when the number of muscle parameters sets are low and number of virtual participants is sufficiently high. For example, in our data set once roughly 40 virtual participants are included, RMS errors are reduced further by increasing the number of muscle parameter sets from 1 to 2 than by increasing the number of virtual participants from 40 to 41. At this point the marginal reductions (or slopes in Fig. 6) are steeper for increases in the number of muscle parameter sets than for increases in the number of virtual participants.

**Figure 6:**
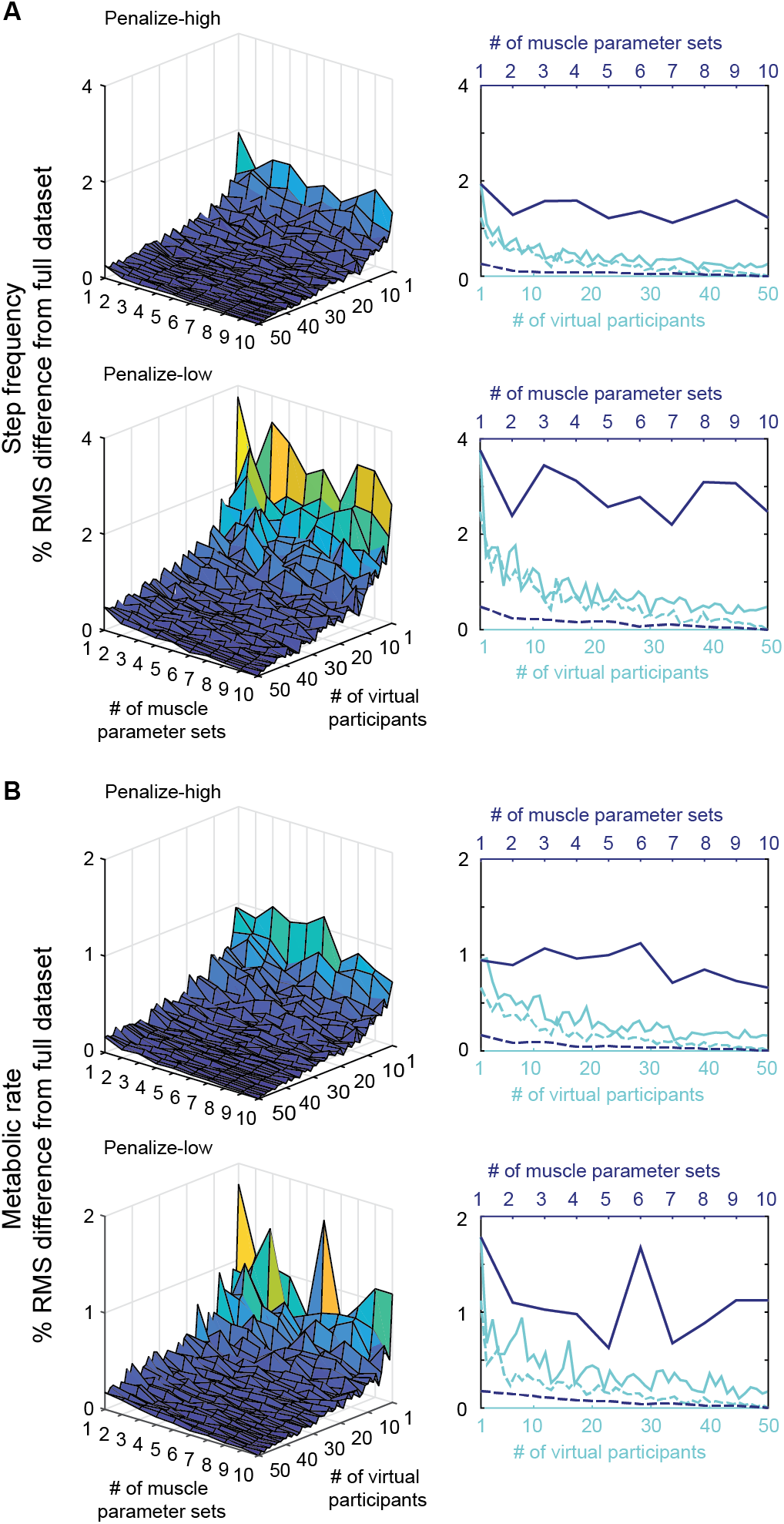
Effect of virtual participant and muscle parameter set sample size. Root mean square (RMS) difference with full dataset as a function of the number of virtual participants and muscle parameter sets for step frequency (A) and metabolic rate (B). Subplots in the right column show cross-sections of the subplots in the left column: with 1 set of muscle parameters (light blue, solid), all sets of muscle parameters (light blue, dashed), 1 virtual participant (dark blue, solid), and all virtual participants (dark blue, dashed).

## 4 Discussion

We used predictive simulations of gait to investigate the effect of an exoskeleton that alters the energetic consequences of walking using a virtual participant design. We were able to generate predictive simulations that well-matched experimental results [4]. While simulated within-stride torques tended to differ from experimental, particularly during stance, our simulated torques averaged across the stride were very similar to experimental. This resulted in full objective, effort term, and metabolic rate landscapes with optima shifted to lower step frequencies under the penalize-high controller condition and higher step frequencies under the penalize-low controller condition, as desired. Our simulated optimal gaits, under each condition, displayed step frequency adaptations consistent with these shifts in optima and indistinguishable from our past experimental results. Simulated individual muscle metabolic rates provided insight, beyond that available from experimental data, into what drives whole-body changes metabolic rate and the new optima. In particular, the slope of the individual muscle energy landscapes change direction under the differing controllers only for muscles that cross the knee. Furthermore, individual muscle stance and swing costs at the optimal gaits reveal distinct coordination strategies consistent with adaptations under each exoskeleton controller condition. Finally, our virtual participant study showed that increasing the number of virtual participants improved simulated outcomes much more than increasing the number of muscle parameter sets.

Our use of predictive simulations is not without its limitations. While trends in metabolic rate and step frequency adaptations were similar between simulation and experiment, magnitudes differed. Across all landscapes, our model predicted metabolic rate was often 0.5-1.5 W/kg higher than that measured experimentally. This is not unexpected. Various metabolic models often produce reliable and reproducible relative changes in metabolic rate, but magnitudes can vary widely between models [40, 41]. In particular, the commonly used Margaria metabolic model we implemented [35] has been shown to produce estimates that tend to be higher than other models [40]. We also found that the step frequency that minimized the full objective under the natural condition was higher than that observed experimentally (2.24 Hz vs. 1.8 Hz, respectively). Interestingly, the step frequencies that minimized effort (2.14 Hz) and metabolic rate (2.16 Hz) alone were more closely aligned with the experimental, indicating that the tracking term in the full objective favored higher step frequencies. This was surprising because the step frequency of the tracked data (1.8 Hz) matched the experimental. Although removing the tracking term entirely would cause gait simulations to become more inaccurate [28], we set the weight of the tracking term to be as small as possible to minimize this effect. We also explored removing the hip angle from the tracked variables (which we expect to be most correlated with step frequency) and including a term to minimize metabolic rate instead of effort in the objective using the model described in [28]. However, in both cases we found that the step frequency that minimized the full objective remained higher than the step frequency that minimized the effort or metabolic rate term.

Simulated individual muscle energetics can offer insight into the full-body energetics and the resulting gait adaptations. First, individual muscle energy landscapes revealed that the slope of the energetic gradient changed direction under the two controller conditions only for muscles that cross the knee. This indicates that the optimal step-frequencies in the penalize-high and penalize-low conditions are largely driven by the energetics of these knee-crossing muscles. These muscles favor a change in step frequency towards a lower resistive torque, while the hip- and ankle-crossing muscles favor a higher or low step frequency independent of the controller condition. However, this is not to say that muscles crossing the hip or ankle do not adapt activity and display changes in metabolic rate. While their slope direction remains largely unchanged, their slope steepness, magnitude and timing of expenditure during gait phases do change under the controller conditions. This indicates that exoskeleton torques are not simply counteracted or offset, but rather the predictive simulations solve for complex and adaptive lower limb coordination strategies. In particular, we observed changes in individual muscle energetics consistent with: i) an adaptation toward higher step frequencies, requiring more work at the hip during swing and relatively less work during a shorter stance phase and ii) an adaptation toward lower step frequencies, requiring more work during push off at the ankle to lengthen steps and relatively less work to swing the limb [42]. Finally, individual muscle metabolic rates may explain why we find a steeper whole-body energetic gradient in the penalize-high condition compared to the penalize-low condition, in both simulation and experiment. It is muscles that cross the knee that drive these steeper slopes; the individual muscle energetic gradients are steeper under the penalize-high than penalize-low condition. This appears to be because under the penalize-high condition, the highest resistive torques are applied during high step frequencies, when knee angular velocity is relatively higher. Conversely, under the penalize-low condition, the highest resistive torques are applied during low step frequencies, when knee angular velocity is relatively lower. High resistive torques applied to muscles moving at higher velocities in turn produce greater muscle work and therefore greater changes in metabolic rate. This leads to a higher cost at +10% step frequency under the penalize-high condition than at −10% step frequency under the penalize-low condition. Another possible contributor is that exoskeleton resistive torques are proportional to knee angular velocity. This occurs in the physical exoskeleton because we used the motor as a generator, where rotational motion induces a voltage in the motor’s windings and in turn a current that generates a magnetic field that resists the motion of the knee [43,44]. At low velocities, current and therefore resistance, cannot be generated. Our simulated controller replicated these effects. Therefore, in both simulation and experiment, lower torques are applied at −10% in the penalize-low condition than at +10% in the penalize-high condition.

Our virtual participant study revealed that adding virtual participants tends to improve simulation outcomes more so than adding muscle parameter. However, at larger numbers of virtual participants, additional muscle parameter sets can meaningfully improve accuracy. If starting with one virtual participant and one muscle parameter set, a 50% reduction in outcome measure RMS error (for example from 4% to 2%) can be achieved by adding roughly 5-10 virtual participants. It is not possible to consistently achieve this same reduction by adding additional muscle parameter sets alone. For most predictive simulation studies, using one muscle parameter set per virtual participant appears to create sufficient variation in the dataset, without adding unnecessary computational cost. However, accuracy can still be improved with a larger muscle parameter set size, especially when the number of virtual participants is large. For example, if 50 virtual participants are included, a 50% reduction in outcome measure RMS error (for example from 0.3% to 0.15%) to can be achieved by adding roughly 1-5 muscle parameter sets. Therefore, in studies where expected differences in outcome measures are on the order of 1% or less, for example when detecting metabolic rate differences between running shoe designs [17], using a larger number of muscle parameter sets could meaningfully improve accuracy.

Our predictive simulation approach has a number of fundamental and applied uses. Fundamentally, it can be used to further investigate aspects of human optimization that are inherently difficult to test experimentally. For example, we can systematically alter the weighting of objectives to predict and potentially understand the effect on gait adaptation. We can also implement physiologically realistic optimization algorithms, such as reinforcement learning, to understand how humans may adapt and learn gaits over time. In a more applied sense, it can be used to explore strategies for improved exoskeleton design and control. While here we used experimental results to validate our predictive simulations, in future we can do the opposite, designing and iterating controllers in advance of human testing to help identify reasonable solution spaces. Our ability to simulate diverse participant pools may also allow us to tailor controller design based on individual users’ abilities or disabilities.

